# Killer Toxins for the Control of Diastatic Yeasts in Brewing

**DOI:** 10.1101/2022.10.17.512474

**Authors:** Victor Zhong, Ximena Garcia, Nicholas Ketchum, Paul A. Rowley

## Abstract

Secondary fermentation in craft beers by diastatic yeasts can result in undesirable consequences, such as off-flavors, increased alcohol content, gushing, and the explosion of packaging. Currently, there are no strategies for inhibiting diastatic yeasts once they have invaded a commercial brewing process. Many strains of yeasts can naturally produce “killer” toxins that inhibit the growth of competing yeasts. Testing the effectiveness of canonical toxins against diastatic yeasts revealed that most (90%) strains are susceptible to the K1 killer toxin. Only two diastatic strains’ resistance to killer toxins was due to toxin production and associated immunity. Four diastatic strains resistant to K1 and K2 toxins were screened against a library of 192 novel killer yeasts to discover novel antifungal activities. This identified novel K2 killer yeasts that were more potent at inhibiting diastatic yeasts than the canonical killer yeast. As proof-of-principle for the control of diastatic yeasts during fermentation, killer yeasts were found to be effective at inhibiting hyperattenuation after a simulated diastatic contamination in a 1,000-liter industrial-style fermentation.

## Introduction

The rise of craft brewing as an industry means that more beer in the marketplace is being produced in facilities lacking pasteurization facilities that stabilize beer against contamination by spoilage yeasts. The shift away from pasteurization is likely because the dominant beer styles produced by this industry are negatively affected by this process. Beer styles that have been aggressively hopped post-boil, such as India Pale Ales (IPAs), will suffer from excessive exposure to oxygen when pasteurized. Therefore, pasteurization can sometimes lead to off-putting flavors described as papery, wet cardboard-like, leathery, or even “catty.” These factors combined mean that even when pasteurization is not prohibited by capital costs, many breweries will opt not to pasteurize as it will cause the degradation of the delicate hop aromas. This aversion to pasteurization can increase the risk of spoilage in craft breweries.

Different species of yeasts are capable of spoiling beer. One particularly problematic group of yeasts are *Saccharomyces cerevisiae* strains that can express the *STA1* gene, producing an extracellular glucoamylase enzyme. These strains have been referred to as diastatic yeasts, a name derived from diastase, an alternative nomenclature for amylase. Diastatic yeasts are an evolutionarily related group of *S. cerevisiae* strains used commercially to produce high-gravity Belgian-style beers (Peter et al., 2018). Diastatic yeasts are unique because the *STA1* gene allows the fermentation of residual polysaccharides such as dextrin and starch (Krogerus and Gibson, 2020). These carbohydrates remain in the beer after the primary fermentation has consumed simple saccharides created in the mashing process. These polysaccharides are usually unavailable to most commercial brewing strains. The *STA1* gene evolved due to a gene fusion between the *FLO11* and *SGA1* (Tamaki, 1978). The fusion resulted in a chimeric protein with the N-terminus of the *FLO11* gene fused to almost the complete open reading frame (ORF) of the *SGA1* glucoamylase. The Sga1 enzyme is involved in the hydrolysis of intracellular glycogen during yeast sporulation. The 5’ end of *FLO11* fused to *SGA1* encodes a signal sequence that enables the transport of Sta1 to the extracellular milieu, where it can hydrolyze residual dextrin and starch to glucose. The released glucose is used to prolong fermentation, referred to as over-super- or hyperattenuation. hyperattenuation results in the overproduction of CO_2_ and alcohol, imparting off flavors and promoting “gushing” and the explosion of packaging. Good hygiene, strain husbandry, and monitoring practices can reduce the likelihood of contamination by diastatic yeasts. Notably, there are no current solutions to prevent diastatic growth after the invasion of a brewing line beyond the destruction of the contaminated product.

Killer yeasts can produce extracellular proteinaceous killer toxins that inhibit the growth of competing species of fungi (Schmitt and Breinig, 2006). Many studies have shown the effectiveness of killer yeasts in preventing spoilage of fruits, silage, and wine fermentation (Jijakli et al., 2002; Kitamoto et al., 1993; Liu and Tsao, 2009; Lowes et al., 2000; Platania et al., 2012; Santos et al., 2011, 2004; Schnürer and Jonsson, 2010). These successes have led to the commercialization of certain fungal species as biological controls in agricultural processes. *Saccharomyces* yeasts were some of the first species identified as producing killer toxins, and surveys have estimated that ∼22% of strains are killer yeasts. Killer toxin expression is often dependent on cytoplasmic double-stranded RNAs (dsRNAs) that are replicated and encapsidated by viruses of the family *Totiviridae* (Schmitt and Breinig, 2006). To date, nine dsRNA-encoded killer toxins are produced by different strains of *S. cerevisiae* (K1, K2, K28, and Klus) and *S. paradoxus* (K62, K1L, K21, K74, K21/K66). Furthermore, at least two functional genome-encoded killer toxins exist in *S. cerevisiae* (KHR and KHS). Each type of killer toxin is distinct and targets susceptible cells by disrupting cell membranes (K1, K1L, and K2) or arresting the cell cycle (K28). Importantly, the antifungal activities of killer toxins are generally limited to closely related species, and there is evidence of widespread resistance to killer toxins across different yeast lineages. Despite this resistance, some studies have shown that the canonical killer toxins of *Saccharomyces* yeasts can successfully inhibit specific human and agricultural pathogens. Moreover, killer yeasts are commonly used in winemaking and could contribute to preventing spoilage by wild yeasts. Importantly, the low pH environment of grape ferments is ideally suited for the optimal activity of all known *Saccharomyces* killer toxins.

In this study, it is demonstrated that the K1 killer toxin from *S. cerevisiae* has potent antifungal activity against the majority of diastatic yeast strains. K1 resistance in diastatic yeast appears not to correlate with K1 toxin production, as diastatic strains tend to produce K2. Intriguing, after screening a large collection of *Saccharomyces* killer yeast, diastatic strains resistant to canonical K1 toxins were found to be susceptible to a non-canonical variant K2 toxin (K2v). As proof of principle for the application of killer yeasts as a control against diastatic contamination, it is shown that a K1 killer yeast strain can prevent super attenuation in an industrial-scale fermentation with no adverse effect on the final gravity of the ferment.

## Results

To determine the susceptibility of diastatic (*STA1*+) strains of *S. cerevisiae* to killer toxins, lawns of cells from 38 diastatic strains were plated on YPD media (pH 4.6) with 0.003% w/v methylene blue. Eight representative strains of *Saccharomyces* yeasts expressing canonical killer toxins (K1, K1L, K2, K21/K66, K28, K62, K74, and Klus) were inoculated onto the seeded lawns of diastatic yeasts. Plates were incubated for three days at room temperature and were checked for the appearance of zones of growth inhibition and methylene blue staining surrounding the killer yeast as an indication of cell death. The growth inhibition of each strain of diastatic yeast was scored according to the degree of growth inhibition (Figure 1). Of all the canonical killer toxins assayed, K1 was judged as being most inhibitory to diastatic yeasts, inhibiting 90% of the strains tested. It was observed that only four diastatic strains (OLY1112, AFA, AQH, and AEA) were resistant to K1.

**Figure 1.**
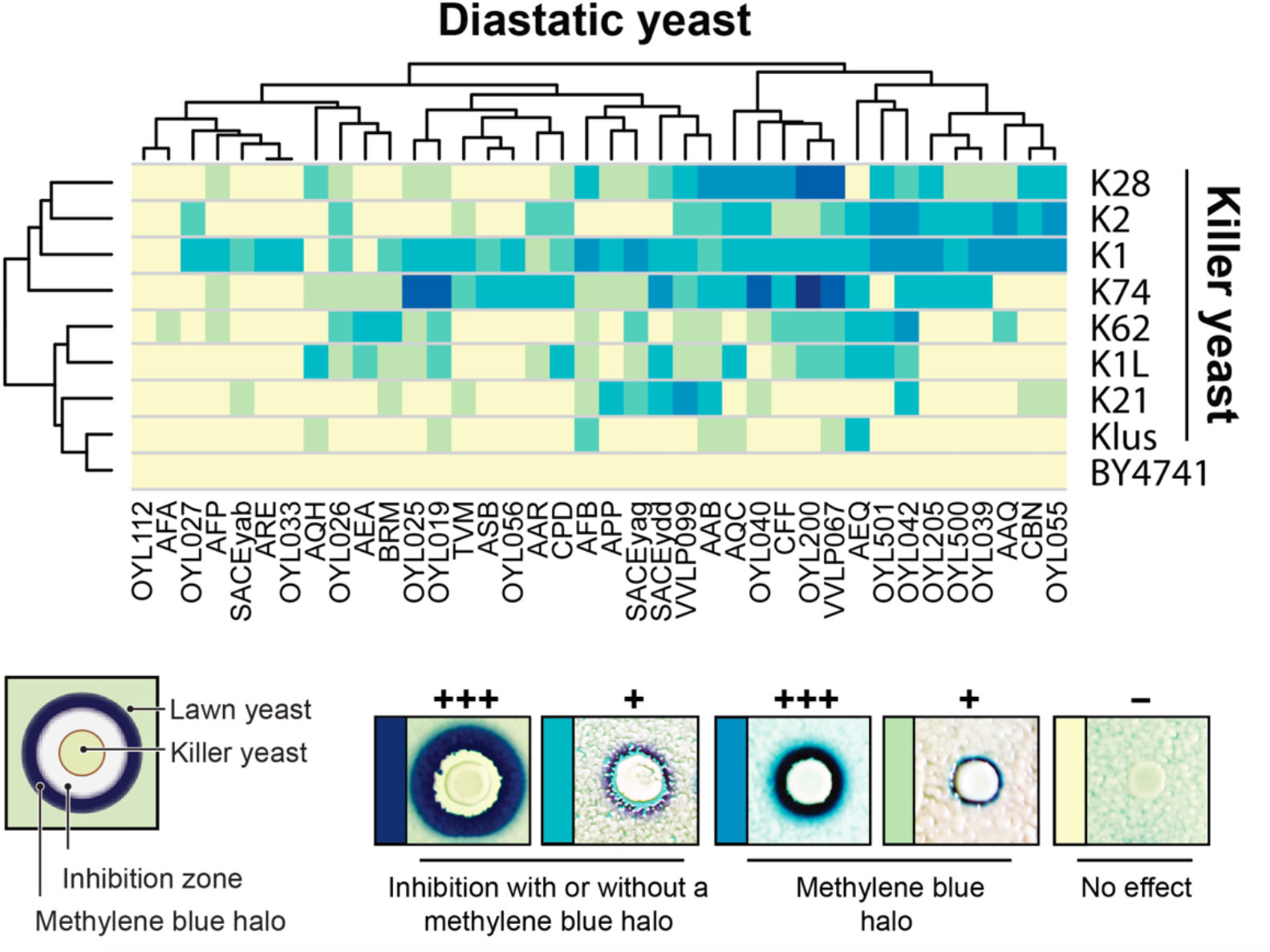
Diastatic yeasts are susceptible to canonical killer toxins produced by *Saccharomyces* yeasts. Killer toxin activity against 38 strains of diastatic yeasts was qualitatively assessed based on the presence and size of zones of growth inhibition and methylene blue staining around killer yeasts as diagrammed (bottom left). Darker colors on the cluster diagram represent a more prominent killer phenotype, with yellow indicating no detectable killer phenotype (bottom right). The non-killer yeast strain *S. cerevisiae* BY4741 was used as a negative control.

Killer toxin production by *Saccharomyces* yeasts is accompanied by immunity to the mature toxin. Thus, killer yeasts are immune to their cognate toxin and other closely related killer toxins. To determine whether the killer toxin-resistant diastatic yeasts had gained immunity due to killer toxin production, each was assayed for their ability to produce active killer toxins. The four K1-resistant diastatic yeasts and an additional 13 strains that were resistant to K2 were used to challenge three lawns of *S. cerevisiae* known to be susceptible to K1 or K2. Only three of 17 diastatic strains were considered killer yeasts (AEA, AQH, and AFB), and all three were able to inhibit the growth of the laboratory strain *S. cerevisiae* BY4741. Importantly, none of the diastatic killer yeasts could inhibit the growth of *S. cerevisiae* CYC1058, a K2 killer yeast. This innate resistance to K2 suggested that these diastatic killer yeast strains were likely themselves K2 killer yeasts. To determine whether killer toxin production in killer toxin-resistant strains was due to viruses and associated dsRNA satellites, nine K1- and K2-resistant strains were subjected to analysis by cellulose chromatography to enrich for dsRNAs (Figure 2A). Agarose gel electrophoresis revealed that three strains contained dsRNAs with sizes the same as totiviruses (∼4.6 kbp), and three strains harbored both totiviruses and dsRNA satellites (∼1.5 kbp) (Figure 2A). No dsRNAs were observed in the three other strains that were assayed. Reverse transcriptase PCR (RT-PCR) indicated that primers specific for the K2 gene could amplify K2 in AEA, AFA, AFB, and AFP, whereas K1-specific primers did not (Figure 2A). Curiously, K2 was amplified from strain AEA, despite the failure to visualize a dsRNA satellite. Exposure to cycloheximide and elevated temperatures was able to cure the dsRNA satellites from AFA, AFB, and AFP. This treatment resulted in the loss of killer toxin production by both AEA and AFB, suggesting that the killer toxin gene was indeed encoded upon a dsRNA satellite in each strain (Figure 2B).

**Figure 2.**
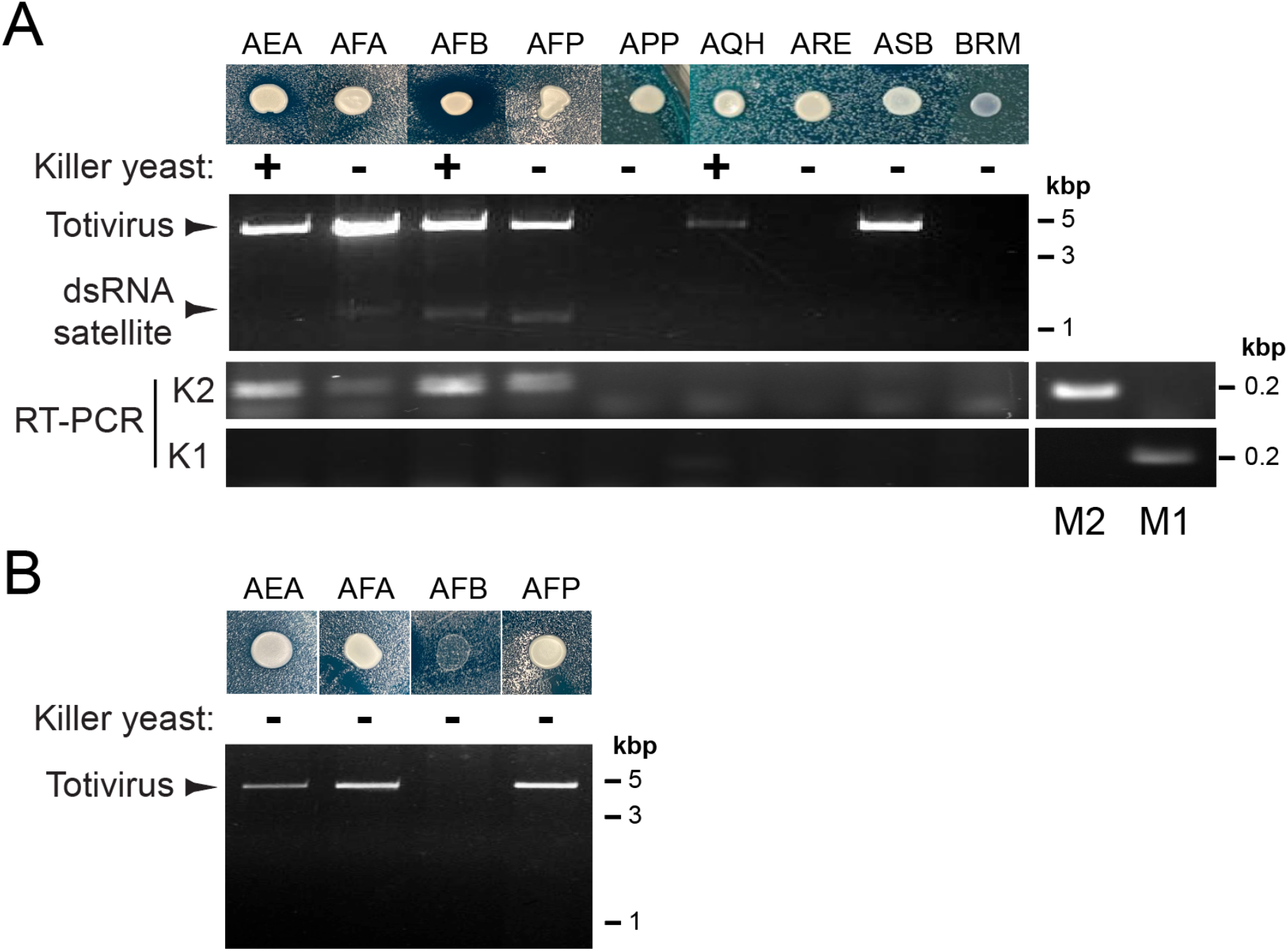
Resistance to K2 is consistent with K2 killer toxin production by diastatic yeasts. (A) The extraction and analysis of dsRNAs from the killer or non-killer diastatic yeasts resistant to K1 and K2 killer toxins. The presence of K1 or K2 genes in purified dsRNAs was probed by rt-PCR and compared to M1, and M2 dsRNAs extracted from K1 and K2 killer yeasts, respectively. (B) Exposure to cycloheximide and elevated temperatures was used to cure diastatic strains of dsRNA satellites and viruses, resulting in the loss of killer toxin production.

Four diastatic yeast strains were resistant to K1 and K2 killer toxins (AEA, AFA, AQH, and OYL112), and only OYL112 appeared to be resistant to all canonical killer toxins. To determine whether there was any *Saccharomyces* killer yeast strain that would inhibit the growth of these yeasts, 192 previously identified and uncharacterized killer yeasts were screened for their ability to inhibit their growth. Of these strains, 67 produced noticeable but sometimes subtle zones of growth inhibition when plated on one of four lawns of K1/K2 resistant diastatic yeast. In particular, the CHD and BSG strains of killer yeasts were able to inhibit all resistant diastatic killer yeasts (Figure 3A). Both of these killer yeasts harbored dsRNAs consistent with the presence of totiviruses and dsRNA satellites and, when probed by RT-PCR, were found to be amplified by primers specific for the K2 killer toxin. This result is of particular surprise as AEA, AFA, AQH, and OYL112 were all found to be resistant to the canonical K2 toxin.

**Figure 3.**
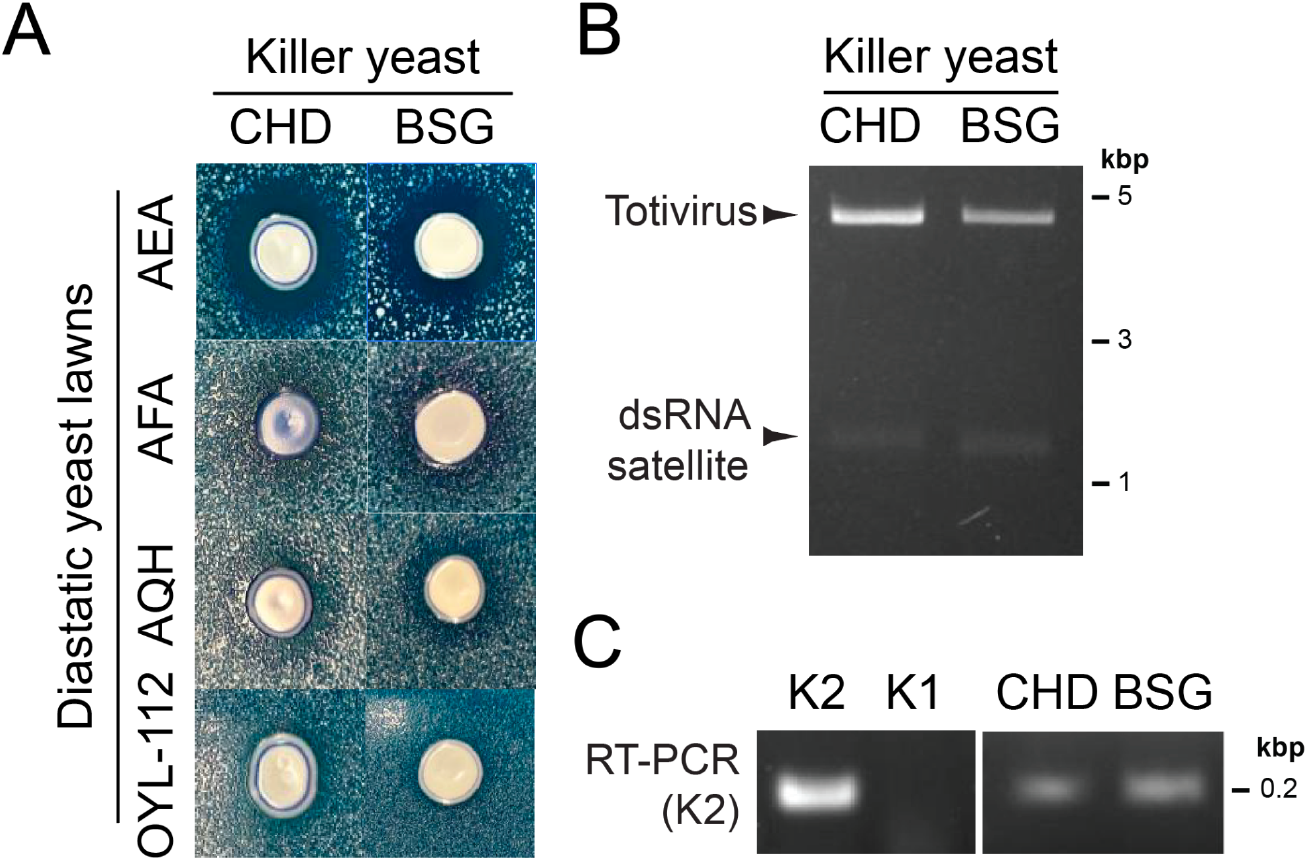
Novel killer yeasts can inhibit the growth of diastatic yeasts resistant to K1 and K2 killer toxins. (A) Agar plate killer assays depict the sensitivity of diastatic yeasts to two uncharacterized killer yeasts. (B) Killer yeasts that inhibit K1 and K2-resistant diastatic yeasts contain dsRNA viruses and satellites. (C) RT-PCR indicates that dsRNAs extracted from strains CHD and BSG are amplified by primers specific to K2.

To determine whether it was possible to prevent hyperattenuation by diastatic yeasts with the addition of killer yeasts, two 1,000-liter brewing trials were conducted. Both fermentation trials proceeded as normal in the first six days, with some variability in the gravity readings in the first 36-hour period due to the rapid evolution of CO_2_. After three consecutive days of stable gravity readings, fermentations were judged to have reached terminal gravity (∼1.6° Plato (P)). In trial one, the diastatic yeast strain Belle Saison (Lallemand Inc.) was added. While the pH remained relatively consistent, the gravity of this fermentation decreased by 0.6° P over ∼7 days to a final reading of 1.06° P before the trial was halted (Figure 4). In trial two, the same diastatic yeast strain was added, but simultaneously with the K2 killer yeast, Viva VIC-23 (Renaissance Yeast). After this addition, the gravity again reduced, but by only 0.08° P. Over the remaining course of the fermentation, the gravity recovered 1.77° P (Figure 4). While it was evident that adding a K2 killer yeast successfully prevented hyperattenuation, there was still a noticeable and undesirable phenolic flavor to the final brew.

**Figure 4.**
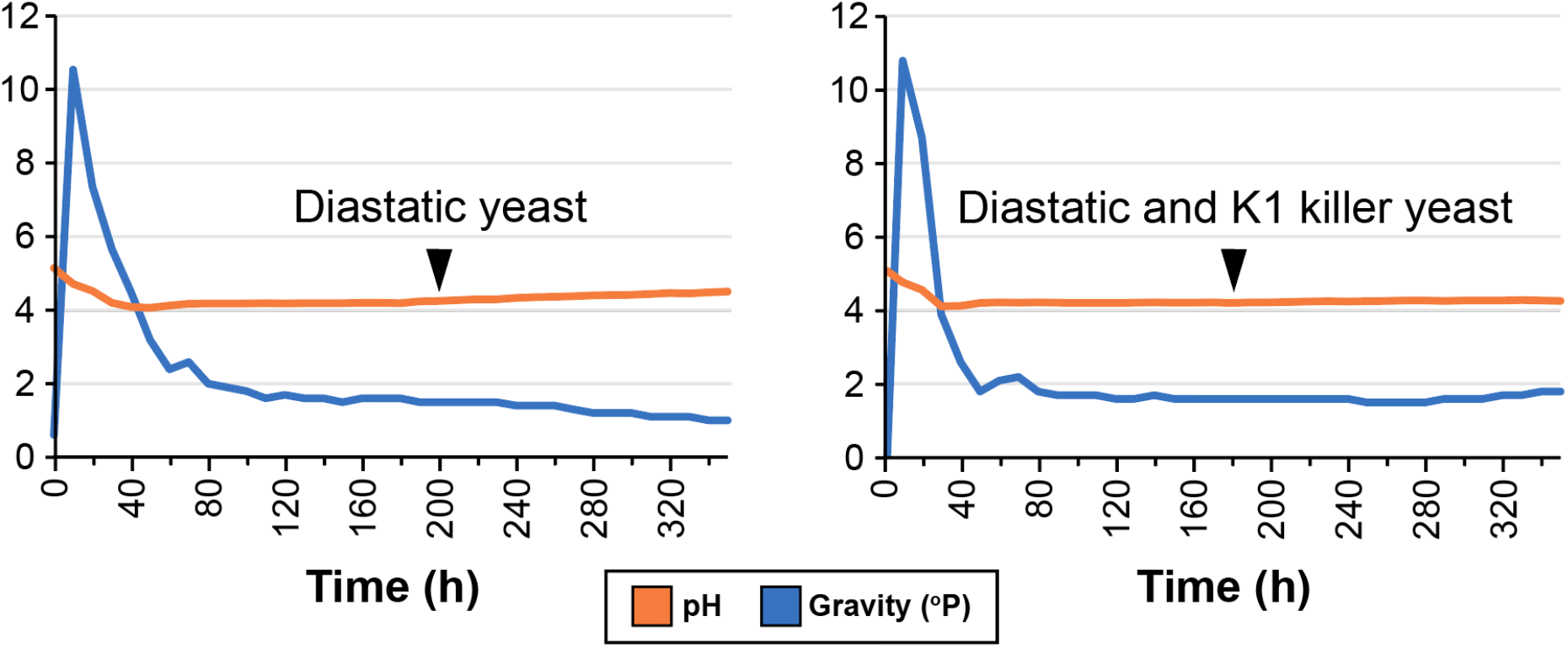
Killer yeasts can prevent hyperattenuation in fermentation trials. The pH and specific gravity of two 1,00 liter scale brewing trials were monitored for 14 days after the addition of diastatic yeast strains after 7-8 days (A) with or without the addition of a K2 killer yeast.

## Discussion

Diastatic contamination of craft breweries is a serious problem for a growing industry that manages this problem by maintaining good hygiene and implementing monitoring practices. In the event of diastatic contamination, the only course of action is the destruction of the contaminated product. In this manuscript, it has been demonstrated that diastatic yeasts are susceptible to proteinaceous killer toxins produced by different strains of *Saccharomyces* yeasts. Furthermore, it is revealed that hyperattenuation caused by contamination by a diastatic yeast strain can be prevented by adding a killer yeast. This discovery offers a unique strategy for controlling and preventing diastatic contamination.

Killer toxins have been known to have a narrow spectrum of antifungal activity that has blunted the enthusiasm for their application against pathogenic and spoilage yeasts. However, numerous laboratory studies have shown the utility of a myriad of killer yeasts in controlling fungal spoilage and the unique susceptibility of certain species of *Candida* yeasts to K1 (Fredericks et al., 2021b). Together these studies have adequately demonstrated the future potential for the application of killer yeasts.

Application of killer yeasts to limit or prevent diastatic contamination could take a variety of forms, such as the addition of a bolus of killer yeast to fermentations upon the detection of diabetic invasion or to engineer or use brewing strains that naturally or are engineered to produce killer toxins. Regardless of their application, one important consideration for the addition of killer yeasts to craft breweries will be the effect on the finished product and to ensure that the addition of killer yeasts or the engineering of brewing strains does not influence desirable flavor profiles. One other consideration would also be that killer yeasts themselves can also invade industrial processes by outcompeting commercial strains of yeasts during fermentation (Maule and Thomas, 1973). However, since killer toxin resistance is widespread amongst commercial brewing strains, using killer toxin-resistant strains would prevent such an invasion.

## Methods

The methods for performing killer yeast assays, dsRNA extractions, and the curing of dsRNAs from yeasts are described previously (Fredericks et al., 2021a).

### Fermentation trials

Trial one was brewed with Rahr two-row brewers’ malt and 0.8 lbs. of bravo hops (20 IBU). The 11.5°P wort was knocked out at 68° F and inoculated with 10 liters of WLP-001 at a pitching rate of 1.0*10^6 cell/ml/°P. Once the gravity had stabilized for three consecutive days, the diastatic yeast was added. 5L of the STA1+ yeast Belle Saison from Lallemand was added through the hop port while CO2 provided positive pressure. Trial two was brewed with Rahr two-row brewers’ malt and 0.8 lbs. of bravo hops (20 IBU). The 11.5°P wort was knocked out at 68° F and inoculated with 10 liters of WLP-001 at a pitching rate of 1.0*10^6 cell/ml/°P. Once the gravity had stabilized for three consecutive days, the STA1+ yeast was added along with the K2 killer yeast. 5L of the STA1+ yeast Belle Saison from Lallemand and 5L of the K2 killer yeast, Viva VIC-23, from Renaissance Yeast was added through the hop port while CO2 provided positive pressure. All data from both trials were collected in real-time via a recirculating inline loop attached via the hop port. The instrument collected data every 30 min on pH, Density (g/cm 3), Gravity (°P), DO (mg/L), Conductivity (uS/cm), temperature (°F), Pressure (PSI) using the BrewMonitor real time data collection system. The instrumentation was cleaned with the alkaline non-caustic CIP cleaner Cell-R-Mastr, triple rinsed with 60°C water, and sanitized with peroxyacetic acid for 30 min before attaching to the fermenter.

## Acknowledgments

We would like to thank the team at Precision Fermentation for their Brew Monitor and all the support we received while performing the trials at the 1,000-liter scale. We would like to thank Laura Burns and Lance Sharner at Omega Yeast for follow-up experiments on their catalog of diastatic yeasts.

